# Murine norovirus allosteric escape mutants mimic gut activation

**DOI:** 10.1101/2025.02.04.636510

**Authors:** Michael B. Sherman, Hong Q. Smith, Faith Cox, Christiane E. Wobus, Gillian C. Lynch, B. Montgomery Pettitt, Thomas J. Smith

## Abstract

Murine norovirus (MNV) undergoes large conformational changes in response to the environment. The T=3 icosahedral capsid is composed of 180 copies of ∼58 kDa VP1 that has N-terminal (N), shell (S) and C-terminal protruding (P) domains. In phosphate buffered saline, the P domains are loosely tethered to the shell and float ∼15Å above the surface. At conditions found in the gut (i.e. low pH with high metal ion and bile salt concentrations) the P domain rotates and drops onto the shell with intra P domain changes that enhance receptor interactions while blocking antibody binding. Two of our monoclonal antibodies (2D3 and 4F9) have broad strain recognition and the only escape mutants, V339I and D348E, are located on the C’D’ loop and ∼20 Å from the epitope. Here we determined the cryo-EM structures of V339I and D348E at neutral pH +/- metal ions and bile salts. These allosteric escape mutants are constitutively in the activated state without the addition of metal ions or bile salts, thus explaining how they escape neutralization. Dynamic simulations of the P domain further suggest that movement of the C’D’ loop may be the rate limiting step in the conformational change and that V339I increases the motion of the A’B’/E’F’ loops compared to wt, making it easier for the virus to transition to the activated state. These findings have important implications for norovirus vaccine design since they uncover a form of the viral capsid that should lend superior immune protection against subsequent challenge by wild type virus.

**Importance:** Immune protection from norovirus infection is notoriously transient in both humans and mice. Our results strongly suggest that this is likely because the ‘activated’ form of the virus found in gut conditions is not recognized by antibodies created in the circulation. By reversibly presenting one structure in the gut and a completely different antigenic structure in circulation, gut tissue can be infected in subsequent challenges, while extra intestinal organs are protected. We find here that allosteric escape mutants to the most broadly neutralizing antibodies thwart recognition by transitioning to the activated state without the need of gut conditions (i.e. bile, low pH, or metal ions). These findings are significant because it is now feasible to present the activated form of the virus to the immune system (for example as a vaccine) to better protect the gut tissue for longer periods of time.

## Introduction

The T=3 icosahedral calicivirus capsid is composed of 180 copies of VP1 with a molecular weight of ∼58 kDa that is divided into the N-terminus (N), the shell (S) and the C-terminal protruding (P) domain (1–5) (Figure 1). The S domain forms a shell around the viral RNA genome, while the P domains form protrusions emanating from the shell comprised of A/B and C/C dimers. The P domain is further subdivided into P1 and P2 subdomains with the latter containing the binding sites for cellular receptors (5–9) and neutralizing antibodies (10–13). For reasons that have not been entirely clear, gut immunity to both human (14) and murine (15, 16) noroviruses quickly wanes and the host can be reinfected by the same strain just months after recovery.

**Figure 1.**
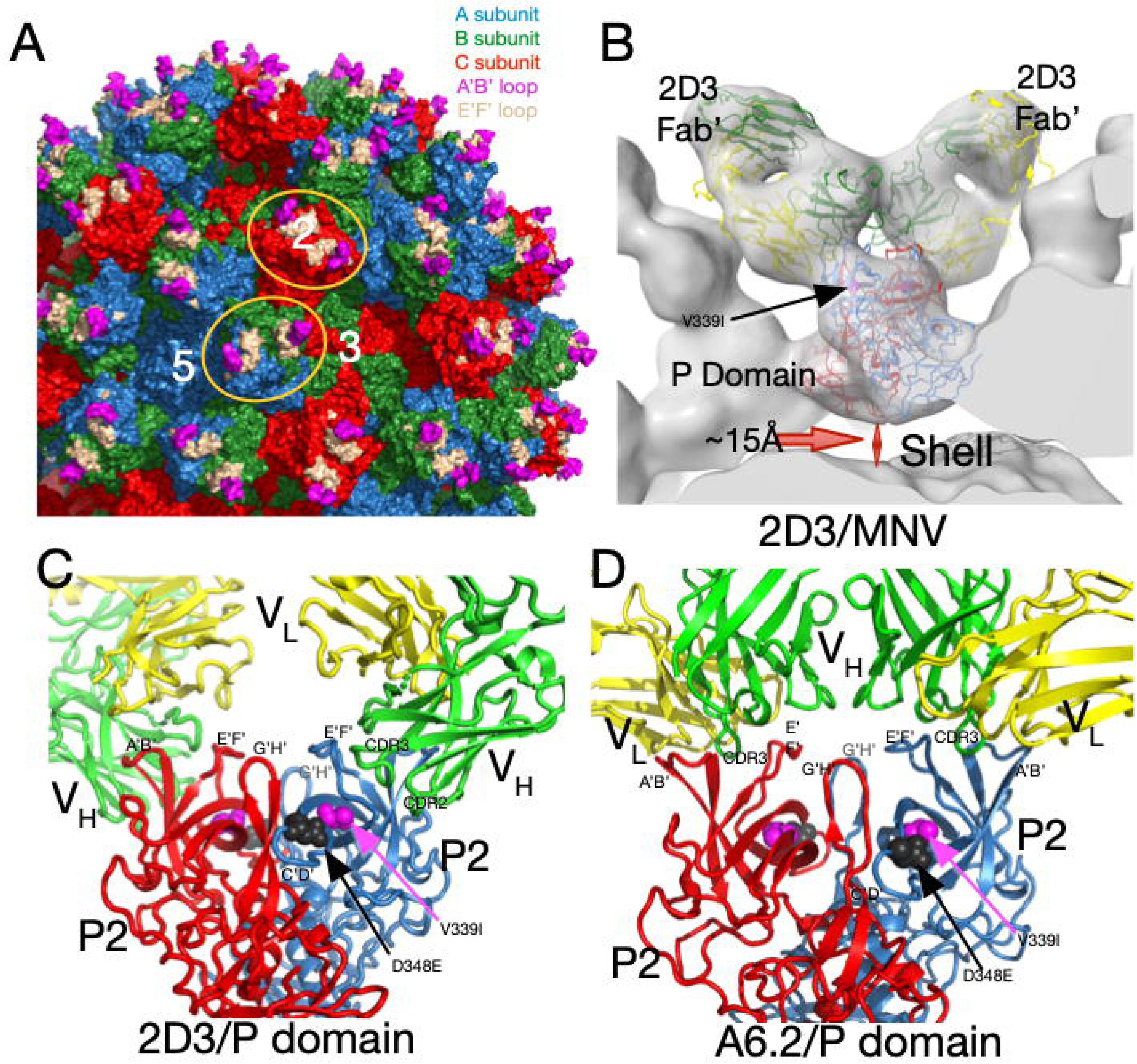
MNV capsid structure and antibody complexes. A) A surface rendered section of the MNV capsid showing VP1-3 colored blue, green and red, respectively. A/B subunits form dimers around the 5-fold axes while the adjacent C/C subunits form dimers at the icosahedral 2-fold axes. Also noted is the location of a 5-, 3-, and 2-fold axis. At the tip of the P domain, the A’B’ and E’F’ antigenic loops are colored mauve and tan, respectively. Example A/B and C/C P domain dimers are highlighted by orange ellipses. B) Previous cryo-EM structure (13) of the 2D3 Fab’/MNV complex and the associated pseudo-atomic model (C). 2D3 binds to the outermost tip of the P domain and makes nearly identical contacts as the antibody A6.2 (panel D (13)). Residue V339I is highlighted in mauve and D348E in black. In panels C and D, the two P domains are colored red and blue and the antibody heavy and light chains are colored green and yellow, respectively. The model from this ∼8Å map only approximates the antibody/MNV complex but is sufficient to show that the 2D3 epitope lies between the A’B/E’F’ loops. D) Shown here is the 3Å cryo-EM structure of the A6.2 Fab/MNV P domain complex. Note that the Fabs in both antibody complexes have nearly the same P domain contacts even though the escape mutations to 2D3 are distal to the bound antibodies and A6.2 escape mutants.

Murine norovirus (MNV, genotype GV.1) is a powerful surrogate for human noroviruses (HNoVs) since it can be grown to high titers in cell culture, has a reverse genetic system available, and mice serve as the natural host as well as a convenient animal model. The latter is crucial since HNoV research will always be limited by the fact that mutants/variants cannot be tested in-vivo. Furthermore, MNV is appropriate as a HNoV surrogate since recent studies on HNoV capsids have recapitulated some of our structural work on MNV (e.g. (17, 18)), and our recent publication (19) showed that the histopathological changes in the mouse MNV model mirror the limited findings in human studies (20–23).

In the original MNV-1 P domain X-ray structure (12), the loops at the tip of the P domain (Figure 1 A) adopted two different conformations; the A’B’ and E’F’ loops were either splayed apart (open) or tightly associated (closed). From our ∼8Å cryo-EM structures of several Fab/MNV-1 (e.g. Figure 1B) complexes (4, 13, 24–26) and the X-ray crystal structure of one of the Fabs (25), we suggested that the antibodies preferred binding to the ‘open’ loop conformation (Figures 1B, C, and D). This was confirmed with our structure of the soluble form of the P domain complexed with the Fab of the neutralizing monoclonal antibody A6.2 to 3.2Å (Figure 1D (27)). As we had predicted, the H chain CDR3 loop ‘unfurls’ to reach down to the hydrophobic cleft between the A’B’/E’F’ loops in the ‘open’ conformation but would not recognize the P domains in the ‘closed’ loop conformation.

We have shown that P domains are flexible and display markedly different conformations depending on environmental conditions. In PBS buffer, the P domains of several caliciviruses (MNV, rabbit hemorrhagic disease virus [RHDV], human norovirus genogroup II [HNoV GII]) are highly mobile and ‘float’ ∼15Å above the surface of the shell (4, 12, 24–26), representing the expanded (apo) form (Figure 2A). Since this structural feature is conserved across calicivirus genotypes and genera (24, 28, 29), we suggested that it likely has an important biological function.

**Figure 2:**
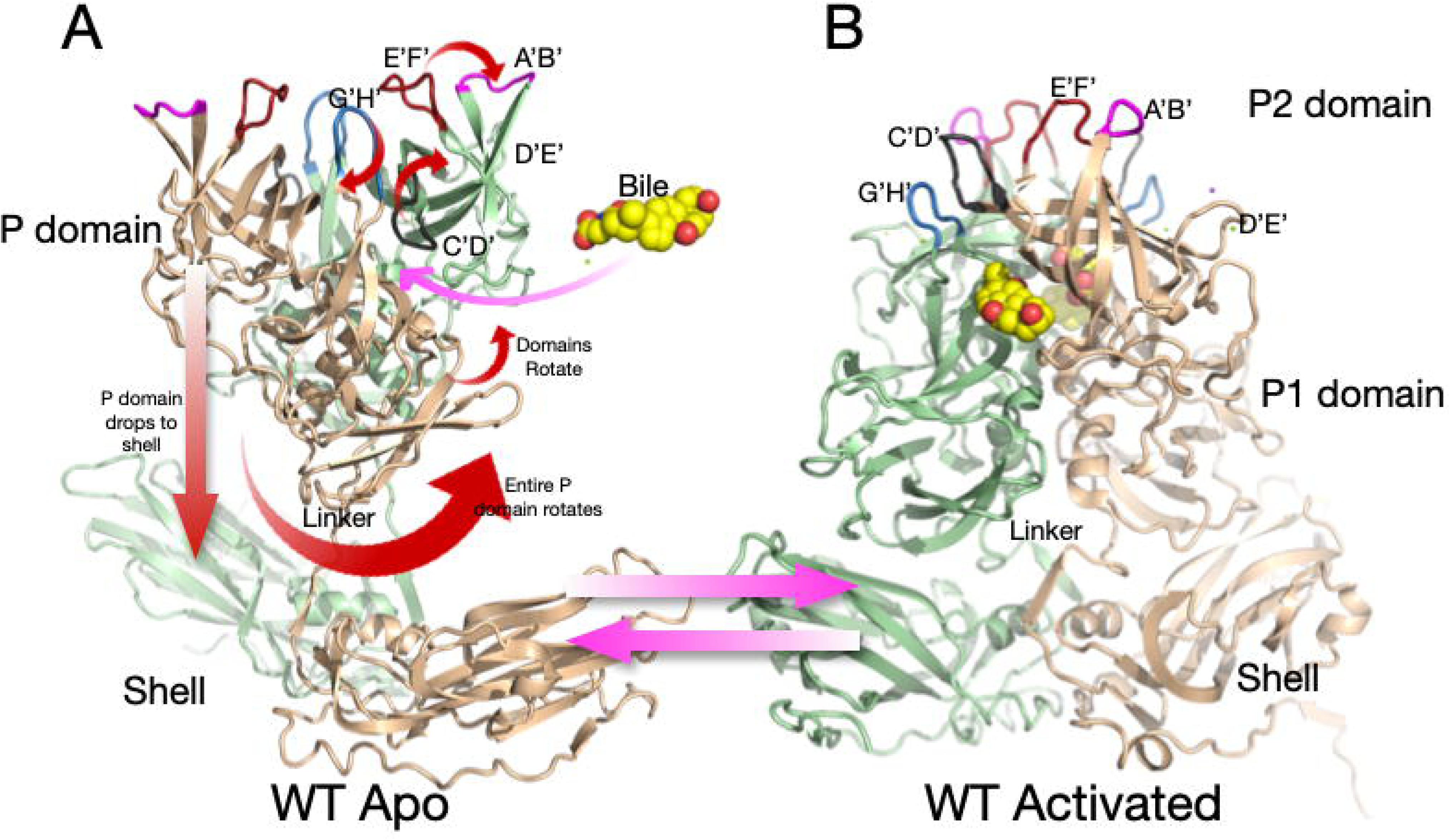
Shown here is the structure of wt MNV-1 in PBS at pH 7.4 (A) and the structural changes that occur during activation (B). In PBS, the P domain floats above the shell and is highly flexible. The addition of metal ions (i.e. Mg^2+^ or Ca^2+^), bile salts, or acidic buffers cause motions noted by the red arrows, resulting in the structure shown in B.

We recently showed that the ‘apo’ structure markedly changes with the addition of the bile acids (e.g. GCDCA) (5), metal ions (30), or low pH (31) (Figure 2B). These changes enhance P domain binding to the receptor CD300lf (8) while blocking the binding of antibodies (27). All these activators cause the same conformational changes (27, 30–32). In the apo form of the capsid, the C’D’ loop points down toward the capsid surface and covers the entrance to the bile binding pocket. This loop orientation allows the A’B’ and E’F’ loops to splay apart into the open conformation. In the presence of bile, low pH, or metals, the C’D’ loop moves up and pushes the A’B’/E’F’ loops into the closed conformation (5). We hypothesize that the G’H’ loop plays a pivotal role in pH- and metal-mediated activation. The G’H’ and C’D’ loops essentially swap spaces at the tip of the P domain; when one loop is up, the other is necessarily down. At neutral pH, a cluster of acidic groups on the G’H’ loop (D440, D443, and E447) is charged and forces the G’H’ loop into a more vertical (radially outwards) orientation allowing the C’D’ loop to fold over the bile binding pocket. At acidic pH or in the presence of metal ions, the charges are neutralized and the G’H’ loop folds down into a compact structure, causing the C’D’ loop to lift and push the A’B’/E’F’ loops into the closed conformation that blocks antibody binding while enhancing receptor binding between the A’B’ and D’E’ loops (8). We also demonstrated that all these activators (i.e., bile acids, metal ions, low pH) cause the P domain dimers to rotate about each other (5, 27, 31, 32) (Figure 2). This rotation changes the surface of the P domain base to complement the shell surface and causes the contraction of the P domain onto the shell. For simplicity, we call the structure in PBS alone ‘apo’ (with open A’B’/E’F’ loops and a floating P domain) while the structure in the presence of metal ions, bile salts, or low pH (with a closed A’B’/E’F’ and contracted P domain) is the ‘activated’ state. It should be noted that while some previous crystal structures of several caliciviruses were in the contracted state (2, 33, 34), our work strongly suggests this was due to the low pH (i.e., pH 4.7-5.6) of the crystallization conditions.

To further explore the dominant epitopes and mechanisms of antibody neutralization, additional antibodies were made against MNV-1 (13, 26). These new monoclonal antibodies, 2D3 and 4F9, broadly neutralize all MNV strains tested (13). Further, a panel of A’B’/E’F’ loop escape mutants to monoclonal antibody A6.2 failed to block 2D3 neutralization (26). The only escape mutants to 2D3 and 4F9 were V339I and D348E (Figure 1 D, E, and F) (13). Since these residues lie on the C’D’ loop and are ∼20Å away from the binding site of A6.2 (26), it was presumed that 2D3 would bind well away from the A6.2 epitope. However, and rather surprisingly, 2D3 binds to the same general location as A6.2 (13). We therefore concluded that these escape mutations act in an ‘allosteric’ manner by indirectly causing conformational changes in the epitope. Herein, we set out to identify the structural changes caused by these escape mutations.

The goal of our studies is to elucidate the mechanisms of V339I and D348E escape and why the 2D3 and 4F9 antibodies broadly neutralize all strains of MNV (26). As reviewed above, at neutral pH and in the absence of metal ions or bile salts (PBS buffer), MNV adopts the ‘apo’ structure where the P domain is ∼15Å off the shell surface and the A’B’/E’F’ loops are in the open conformation (Figure 2A). However, we show here that the allosteric escape mutants, V339I and D348E, are in the ‘activated’ state in PBS where the P domain is tightly associated with the shell and the A’B’/E’F’ loops are closed and thus sterically block antibodies from binding (Figure 2B). These loops are less ordered than when metal ions or bile salts are added, but the escape mutants nearly fully activate the virus without stimuli. Further, while these escape mutants arose using the 2D3 antibody, they also block neutralization by the A6.2 antibody. The reason we did not previously isolate V339I and D348E escapes among the A6.2 escapees is likely because the other mutations in the E’F’ loop were sufficient to block binding and more fit than the V339I and D348E mutants. As shown in figure 2, this transition between apo and activated structures involves many steps and it has been unclear as to the order or importance of each. To better understand this transformation and its regulation, all-atom molecular dynamics simulations were performed starting from the crystal structure of the apo form (12) with and without the V339I mutation. This starting structure presents both the ‘open’ and ‘closed’ conformations of the A’B’/E’F’ loops in the dimer, with the C’D’ loop in the down position. While the A’B’/E’F’/D’E’ loops were highly mobile during the simulations, the C’D’ loop remained intransigent, suggesting that its movement is far slower than the other, less constrained, loops at the tip. The V339I P1 domain was far less mobile than the P2 domain, but the two domains in the dimer rotated about each other akin to what happens during activation (27, 30, 31, 35). Finally, the A’B’/E’F’ loops were more mobile in the V339I than wild-type (wt) suggesting that the mutation increases the dynamic nature of the outer loops and perhaps this facilitates the transition to the ‘closed’ state in the absence of activators.

In summary, this highly mobile virus not only leverages conditions in the gut lumen to block antibody binding but can also trigger these changes via allosteric escape mutations. These results may offer a possible explanation for the transient immune response to norovirus infection and strongly imply that vaccines can be greatly improved by using the activated structure found in gut conditions as antigen.

## Results

### Antibody neutralization

We have previously isolated and characterized three neutralizing monoclonal antibodies (mAb) to MNV-1 called A6.2, 2D3, and 4F9 (4, 12, 13, 25–27, 31) with different neutralization properties and non-overlapping escape mutations. The first mAb studied, A6.2 (36), is an IgG antibody that binds between the A’B’/E’F’ loops (Figure 1D; (27)). A naturally occurring escape mutation was found in the E’F’ loop at residue L386 (37) and subsequent studies identified additional escape mutations in the E’F’ loop at amino acid positions V378, A382, and D385 (25). All these residues were either in contact with bound A6.2 or immediately adjacent. Subsequently isolated antibodies, 2D3 and 4F9, are both IgA antibodies and neutralized the A6.2 E’F’ loop escape mutants (26). The only naturally occurring escape mutations to these antibodies were in the G’H’ loop (D348E and V339I) that do not contact the bound antibody (13).

While we previously showed that 4F9 and 2D3 neutralize viruses with the A6.2 escape mutations on the E’F’ loop (26), we did not test whether mutant viruses with amino acid changes D348E and V339I are neutralized by A6.2. As shown in Figure 3, both mutants are very effective at escaping neutralization by all antibodies. Therefore, whatever conformational changes are induced by the V339I mutation, it disrupts the epitopes for all three antibodies. It is interesting and relevant to note that while A6.2 recognized the MNV capsid protein in Western blots (25), neither 4F9 nor 2D3 can (26). It may be that A6.2 recognizes the outermost portions of the A’B’/E’F’ loops that are better reconstituted in a Western blot while 2D3 and 4F9 are necessarily interacting with more of the 3D structure at the tip. This suggests that while all three antibodies bind to approximately the same region of the P domain (13), they differ in which residues are critical for binding.

**Figure 3:**
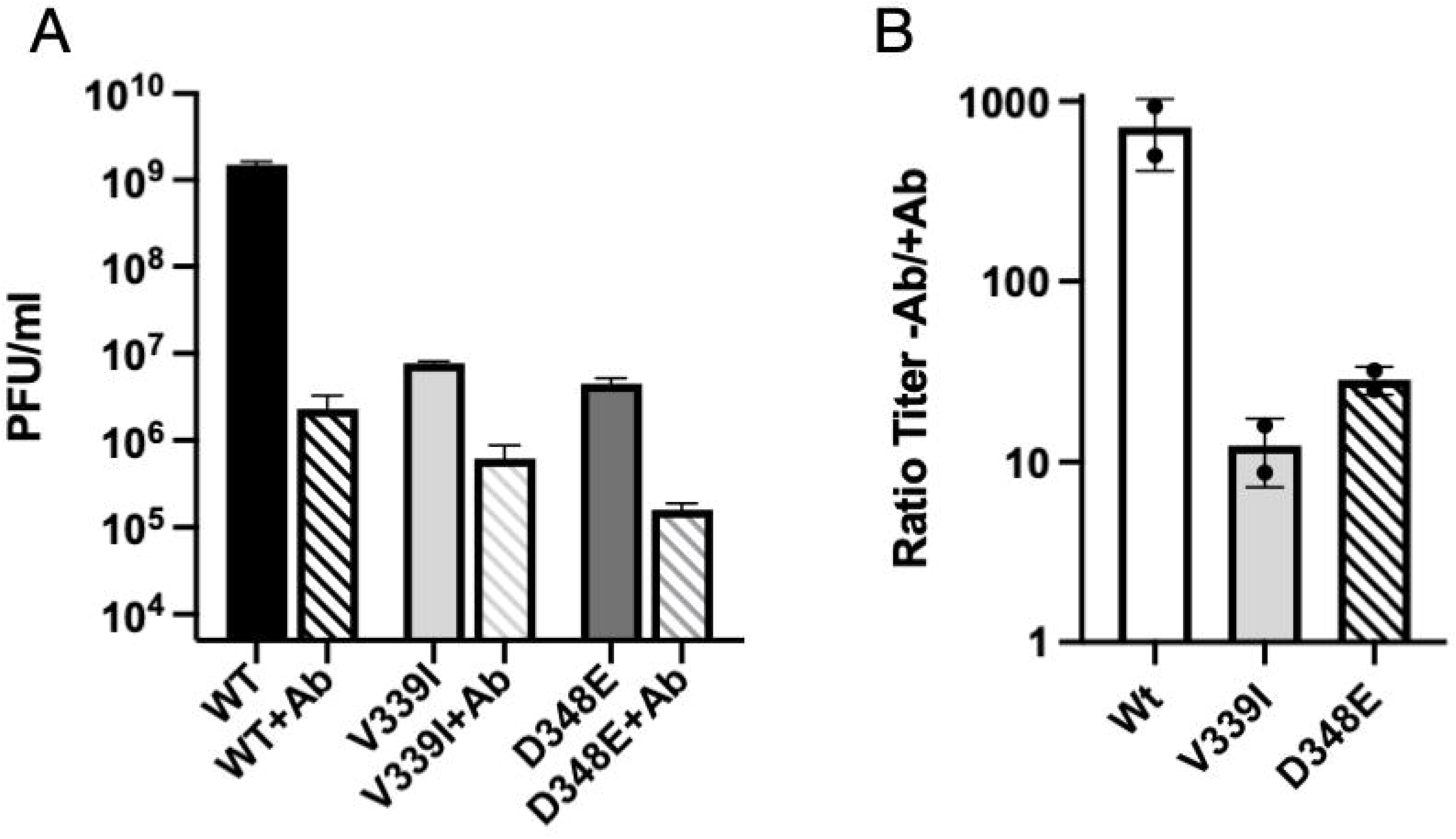
A6.2 neutralization of wt and the V339I and D348E escape mutants. A) The titer of the three viruses in the presence and absence of 50µg/ml monoclonal antibody A6.2. B) Since the viruses all had different initial titers, the ordinate axis in this graph shows the titer in the absence of antibody divided by the titer in the presence of antibody. As shown here, wt was neutralized by more than 3 logs while both mutants were only neutralized by one log.

### Structure of V339I +/- activators

To understand the mechanism of antibody escape, we pursued structural studies of the escape viruses V339I and D348E starting with V339I. Once the sequence of the purified mutant virus had been verified to contain the V339I mutation, the structures of V339I apo, V339I + 2mM CaCl_2_, and V339I + 10 mM GCDCA were determined from that same sample. The calculated resolutions for all three cryo-EM reconstructions were ∼2.7Å. The most striking finding was that, unlike apo wt virus in PBS (Figure 2A), apo V339I was in the activated state with the P domain rotated and lying on the surface of the shell (Figure 4A). As evidence that this is truly V339I without activating ligands, the inset in Figure 4A shows the absence of density where GCDCA would be located. The density of the outermost loops (mauve arrow, Figure 4A) is disordered and for these regions, the unsharpened reconstructions were used to better place the main chain atoms. Figure 4B shows the structure of V339I+GCDCA. There is clear density for GCDCA (inset) and the loops at the tip are more ordered than with apo V339I. Similarly, V339I+Ca^2+^ is in the fully activated state with the P2 domain loops and bound calcium is clearly defined (Figure 4C). The inset shows the density of the G’H’ loop with a bound metal calcium ion. Therefore, the V339I mutation is sufficient to convert the virus to the activated state at neutral pH even in the absence of metals and bile salts, but adding activators took the virus further into the activated state by stabilizing the outer loop conformations. We have previously shown that neutralization of MNV by A6.2, 2D3, and 4F9 are all blocked when the virus is in the activated state (5, 27, 31, 32). Therefore, this mutation driven activation process is a simple answer as to how V339I can allosterically block neutralization by all three antibodies ((26) Figure 3).

**Figure 4.**
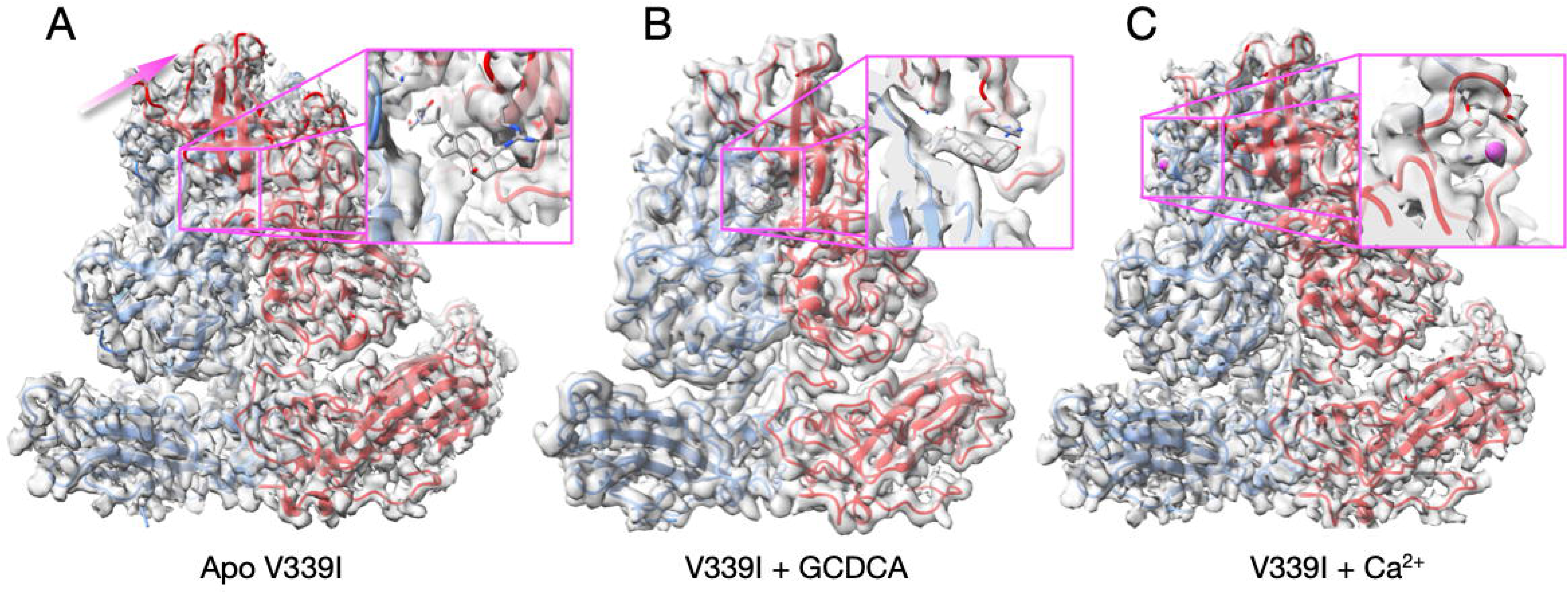
Cryo-EM structures of the V339I mutant virus apo (A) or complexed with GCDCA (B) or calcium (C). A) In contrast to apo wt virus (Figure 2A), the P domain is contracted onto the shell even in the absence of bile salts or metal ions. The one notable difference is that the outermost loops in apo V339I (panel A, mauve arrow) are less ordered than in the presence of either GCDCA or Ca^2+^. The inset figure shows where GCDCA would bind if it were present. The lack of density demonstrates this was indeed an apo sample. B) The V339I+GCDCA structure is also in the activated state and the inset shows clear density for the bound bile salt. The outermost loops are better ordered than in the apo form. C) The density and structure of V339I+Ca^2+^. As with the GCDCA complex, the outer loops are better ordered. The inset shows calcium bound to the G’H’ loop and was not observed in either the bile bound or apo forms of V339I.

Comparisons of the structures of V339I +/- activators is shown in Figure 5. It should be noted that this is the first time we could compare MNV virus structures in the same conformation +/- activators. Panel A shows an overlay of V339I apo (blue and green) onto V339I+GCDCA (red and orange). The positions of the P domains are essentially identical and are in the activated state, resting on the shell surface. The overall RMSD of the alignment between these two structures is ∼0.4Å, well within the margin of error at this resolution. The only apparent differences between the two structures are in the outermost loops that are more than likely due to inherent flexibility. To show the regions of highest disorder, the ribbon diagrams in Figure 5, panels B and C are colored red to blue as the correlation coefficients (CC) of the residues extends from 0 to 1.0. These outermost loops become more ordered with the addition of GCDCA or Ca^2+^ (the color shifts from reds/yellow to blues/greens) that is likely due to stabilized positions of the G’H’/C’D’ loops (insets in Figures 5 A-C).

**Figure 5.**
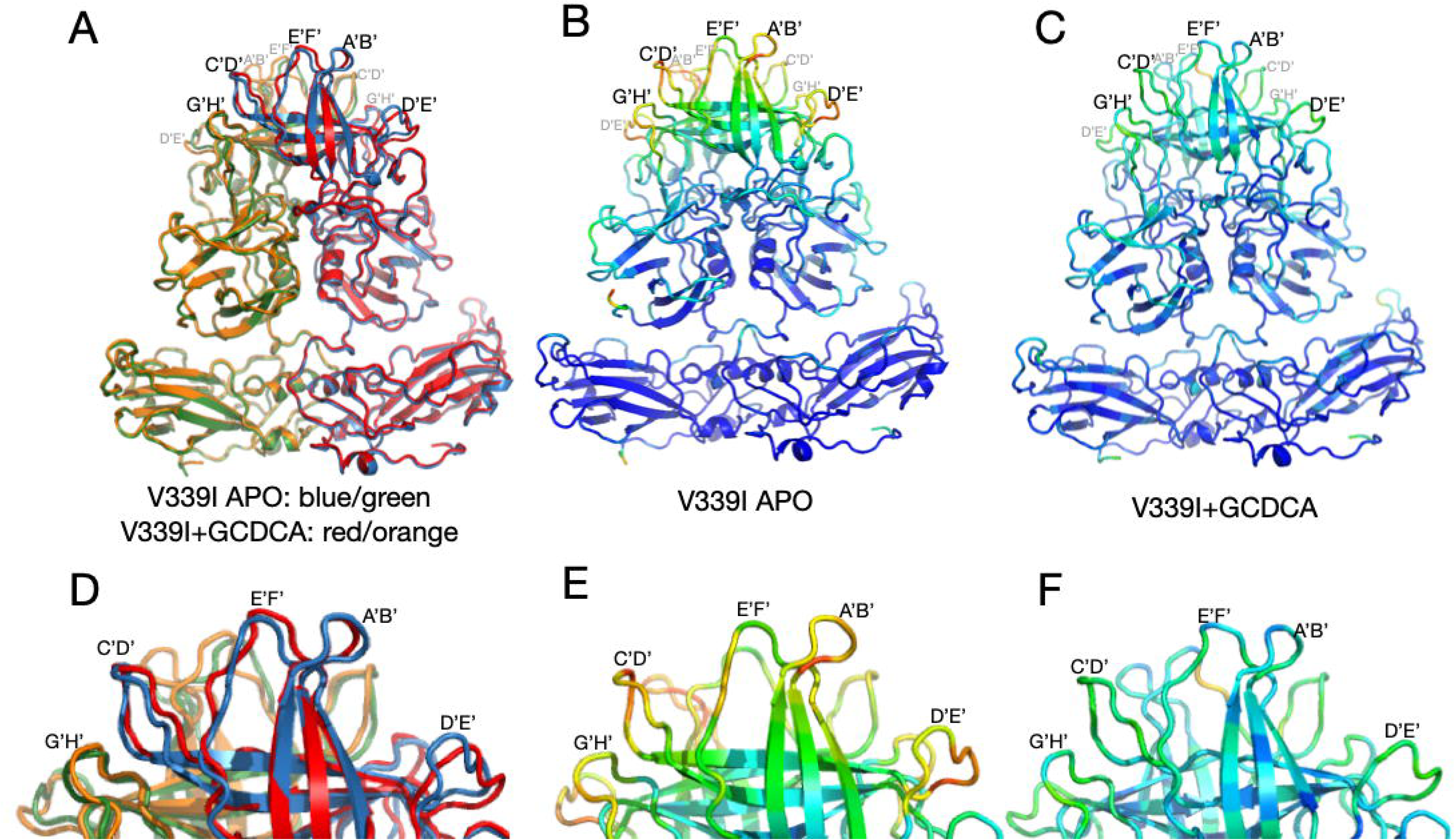
Comparisons of the various V339I structures. A) Overlay of the apo V339I structure (blue/green) onto the V339I+GCDCA structure (red/orange). While the shell and the P1 domains are essentially identical, the structures of the loops at the tip of the P2 domain diverge with the density of the loops in the apo structure being weaker than in the bile bound structure. B,C) Shown here is the per-residue CC for the bile bound (B) and apo (C) mapped onto the structure as B-factors using the formula 100(1-CC). While shell and P1 domains for both structures match their respective densities very well, the outer loops of the apo structure are not well defined (red and yellow colors). Panels D, E, and F are enlarged images of the loops in A, B, and C, respectively. Note that bile stabilizes the conformations of the outer loops as per the shift from red/yellow to blue/green.

These changes in loop flexibility are further shown in Figure 6. In panel 6A, the per residue CC for the C subunits are plotted for the three structures. The overall CC for the P domains alone were 0.69, 0.82, and 0.76 for the apo, +Ca^2+^, and +GCDCA, respectively. The S subunits for all three structures had the highest CC and therefore were the least mobile portions of the capsid. Slightly more mobile than the S domains were the P1 domains (denoted by the orange bars above the graphs). The P2 domain was the most flexible part of the P domain with the outermost loops having extremely weak density in places. The addition of activators (i.e. GCDCA or CaCl2+) decreased this P2 domain mobility as per the improved CC. Therefore, while the V339I mutation alone is sufficient to push the P domain nearly into the fully activated conformation, the addition of these activators finishes the activation process and stabilize these key loops. We had previously noted that there are also differences between the A/B and C/C dimers (5). These dimers differ in their interactions with the shell and with other dimers. The overall CC for the V339I apo P domains of the A and C subunits are 0.69 and 0.67, respectively. While this is a relatively small difference, the densities for the loops and the carboxyl end of the protein appear to be consistently weaker (Figure 6B). This suggests that the context of the P domain within the icosahedron may also influence the mobility of these loops.

**Figure 6.**
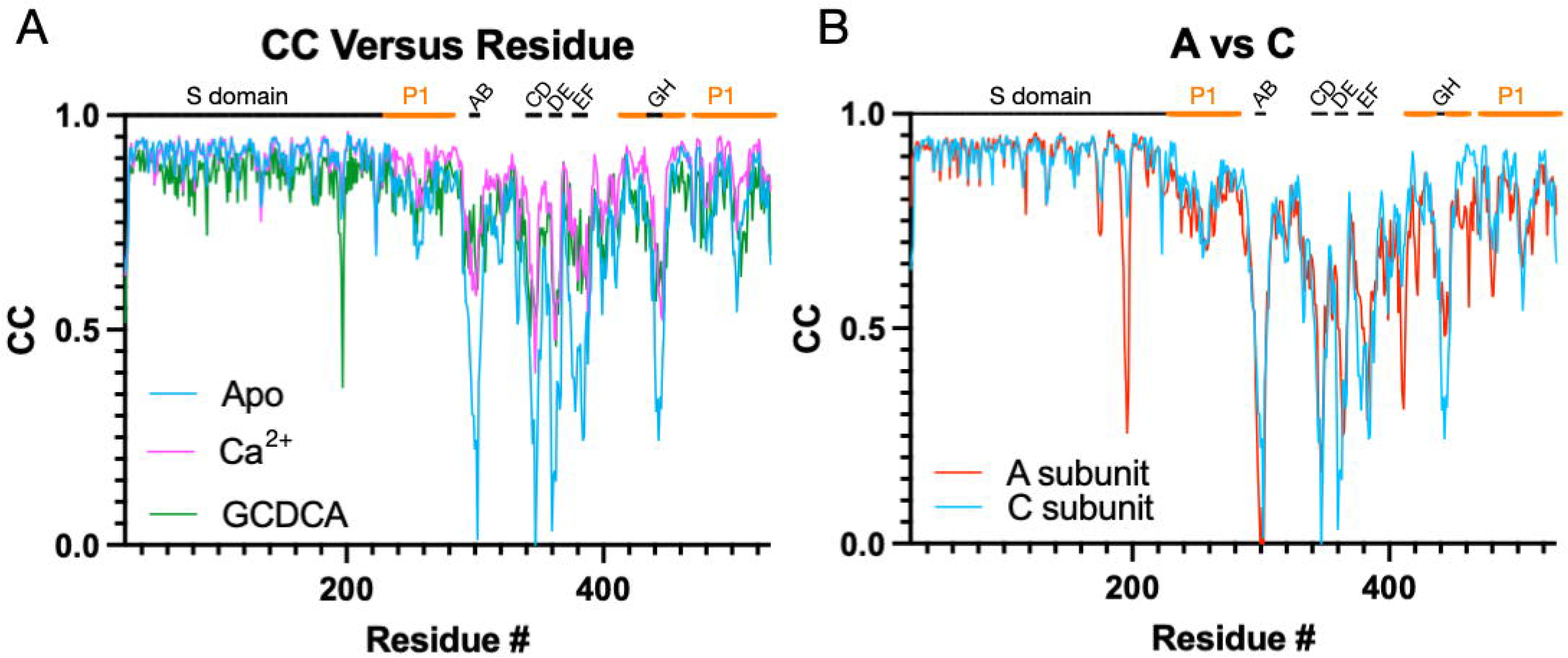
Comparisons of the per residue correlation coefficients among the three structures. A) Shown here are the per-residue CC for the C subunits in the presence and absence of bile or calcium ions. The locations of the outermost loops are noted at the top of the graph and the P1 domain is noted by the orange bars. Although the apo form is in the activated state, the loops are significantly less ordered than when bile or calcium is present. B) Shown here are the per-residue CC of the A versus C subunits in the apo V330I structure. While both structures have highly mobile loops, the A subunit appears less ordered in several loops and from residue 400 onward.

Using the structures of V339I in the activated state and apo wt that is recognized by antibodies (27), we can hypothesize how this allosteric escape mutant causes activation of MNV. Figure 7 shows the apo structure of wt MNV (red and blue) aligned with the activated structure of the V339I mutant (pink and light blue). Highlighted in yellow are the sidechains of the apo wt structure in the immediate vicinity. Overlaid onto the V339 sidechain is a simple substitution with an isoleucine (red). In the apo wt structure, the environment of the V339 is markedly hydrophobic and tightly packed. There is simply not enough space in this cavity to accommodate the extra methyl group of Ile that would clash with I337, F307, and F355. We propose that the mutation causes a shift up in the C’ and D’ strands that results in the upward C’D’ loop movement. Therefore, this overcrowding mutation is sufficient to push the conformation of the C’D’ loop into the ‘up’ position, leading to the cascade of other conformational changes that activate the virus (Figure 2).

**Figure 7.**
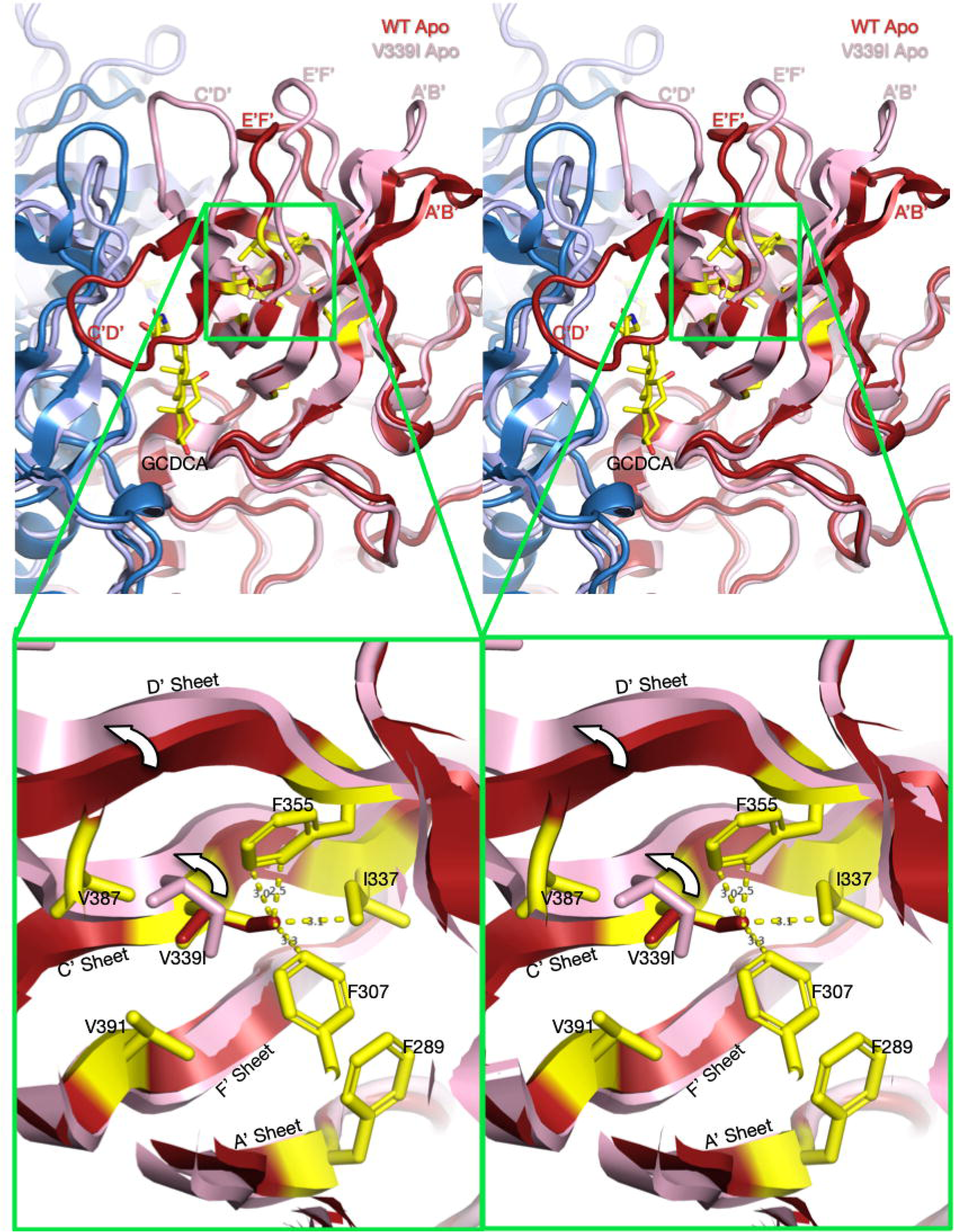
Stereo diagrams of the area surrounding the V339I mutation and possible mechanism for activation. The top stereo pair shows a wt apo P domain dimer (4, 24) in deep red and blue. Overlaid on that structure is the apo V339I structure in pink and pale blue. To help define the location and orientation of the figure, the bound GCDCA from the V339I/GCDCA structure has been added. Note that V339I lies at the N-terminal side of the C’D’ loop that moves up drastically in the wt virus at low pH or when metal ions or bile salts are added. In the magnified view, V339 was replaced with an Ile (red) in the apo wt structure. As noted in this figure, the extra methyl group of the isoleucine would be too close to F335, I337, and F307 in this tightly packed hydrophobic pocket. However, in the actual apo V339I structure, the C’ and D’ β-strands move up, away from the core and make room for the larger isoleucine sidechain.

### Structure of apo D348E

To understand whether V339I and D348E escape mutants use the same mechanism of escape, we next determined the structure of the D348E virus. D348E is on exposed end of the C’D’ loop, far removed from the antibody binding site (Figure 1 C and D). Since D348E can escape neutralization by all three monoclonal antibodies, it was expected that, like V339I, the apo structure would also be in the activated conformation in PBS. As with the V339I mutant, the sample used for structural studies was sequenced immediately prior to cryo-EM data collection and verified to contain the mutation. Figure 8A and B shows that indeed, apo D348E is in the activated state where the P domain is resting on the shell. The loops in apo D348E are in the same conformation as wt under activating conditions (Figure 8C), but far more disordered (Figure 8D). Therefore, the D348E allosteric escape mutation activates MNV in the same manner as V339I.

**Figure 8.**
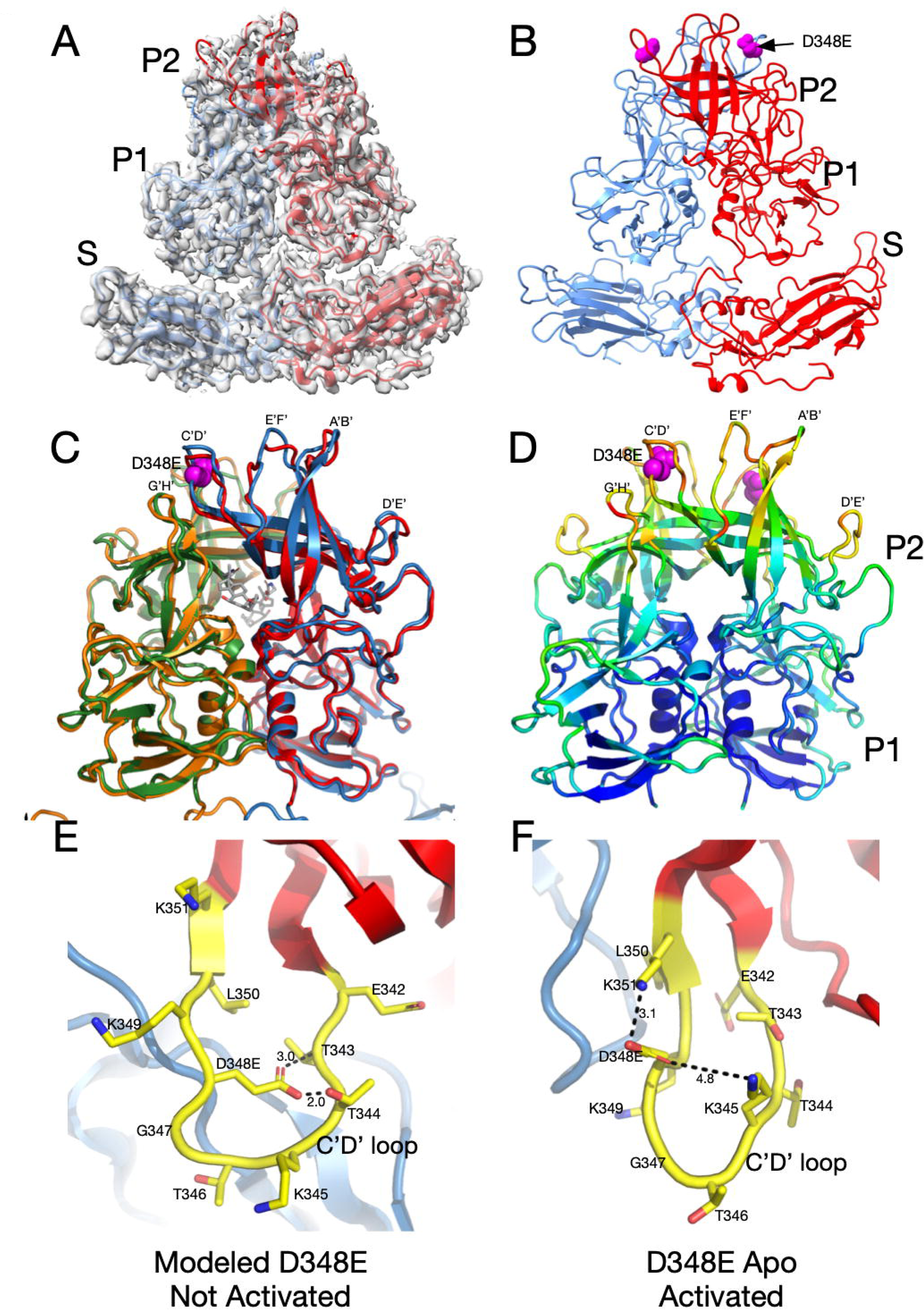
Structure of the D348E allosteric escape mutant and possible mechanism of activation. A) The 3Å EM density of apo D348E and the corresponding model. The two subunits in the dimer are colored red and blue. Note that, as with V339I, the P domain of apo D348E is resting on the shell in the activated conformation. B) Shown here is the apo D348E structure alone with the location of the mutation site denoted by the mauve spheres. C) An overlay of the apo D348E structure (blue/green) with the cryo-EM structure of wild type with GCDCA bound (red/orange). The location of the D348E mutation is noted by the mauve spheres and the bound GCDCA (wild type structure) is represented by the grey model. D) This is the P domain portion of the apo D348E structure colored according to the CC, ranging from blue (1.0) to red (0.0). E) Shown here is a modeled structure of D348E in the unactivated conformation. As with V339I, since the D348E mutation forces the virus into the activated conformation, the mutation needed to be modeled into wt apo conformation. D348 was mutated in COOT and assigned the most likely rotamer position. The distances in this hypothetical model are only shown for reference. The larger sidechain collides with the T343 and T344 and therefore the mutant C’D’ loop cannot adopt this conformation. D) Shown here is the actual structure of Apo D348E that is in the activated conformation where the C’D’ loop is lifted away from the shell. This places D348E out of the plane of the loop and it extends upward where there is sufficient space and places D348E into a favorable electrostatic environment adjacent to K351 and K345.

We proposed above that the V339I mutation pushes the P domain to the activated state by being too large for the space occupied by the C’D’ strands and causing movement in the C’D’ loop and follow-on domino effects throughout the P domain. Since the P domain in the D348E mutant is in the activated conformation without activators, we can only model what may happen when the aspartate is mutated to glutamate in the non-activated, wt apo state (Figure 8E). The side chain of D348 points towards the N-terminal side of the C’D’ loop. When the aspartate is replaced with a glutamate in the modeled structure, it is too large and collides with T343 and T344. However, in the activated conformation of apo D348E, the D348E sidechain rotates up and there is more than enough space to accommodate the larger sidechain. In addition, this orientation places the acidic sidechain in an electrostatically favorable location, adjacent to K351 and K345. Therefore, D348E could cause a push/pull structural change where the clashes caused by the mutation pushes the C’D’ loop out of the wt apo conformation while simultaneously being pulled towards the activated state conformation because of favorable electrostatic interactions with adjacent lysine residues. Movement in the C’D’ loop is then anticipated to cause a domino effect in the P domain leading to closed A’B’ and E’F’ loops at the tip of the P domain, a structural conformation which is resistant to neutralization by all three mAbs.

### Molecular Dynamics Simulations

What we have shown so far is only the static apo and activated structures of MNV, leaving much unknown about the transitional states that conforms one structure into the other. To consider this conformational transition and how V339I might affect it, a series of molecular dynamic simulations were performed. In these two simulations, we used unbiased all-atom simulations to study the effect of the effect of the V339I mutation on the conformational stability of the P domain of MNV and to better understand how this mutation hinders antibody binding. The simulations started from the crystal structure of the apo P domain (12) that has chain A in the open conformation, chain B in the closed conformation, and the C’D’ loop pointing down away from the top. Interestingly, this crystal structure suggests that, with the C’D’ loop pointing down, the A’B’/E’F’ loops can adopt both conformations. With the two subunits in ‘opposite’ conformations, the simulations could show changes in the apo conformation of the V339I mutant using the same model and in the same computation.

The radius of gyration (Rg) is a measurement of the size and compactness of the structure, and values were calculated using all non-hydrogen atoms. As shown in figure S1, both wt and V339I remain stable during the simulations and well converged. However, V339I mutant possibly exhibited slightly more fluctuations as per greater deviations from the mean radius.

The conformational changes were sampled over 900 ns, after 5 ns of equilibration. For the root-mean-square deviation (RMSD) calculations, each snapshot was first aligned to the first frame after equilibration using the non-hydrogen atoms of the backbone of the two subdomains, P1 and P2. The RMSD reflects the room temperature flexibility of the MNV and, specifically, of the P2 domain where the A’B’ and E’F’ loops fluctuate. As shown in Figure S2, there is a marked difference between the P1 and P2 domains where the P2 domain exhibited far more conformational movement than the P1. The P1 domain structure converged quickly and exhibited fluctuations of ∼1.3Å. In contrast, the root-mean-square deviation of the P2 domain varied between 2 and 3 Å, with occasional, short lived, sampling of conformations with RMSD values between 3 and 3.5 Å. The system was apparently converged after 200 ns with respect to that reference frame but not to one single state. These findings that the inhomogeneous fluctuations in the P2 domain are much larger than the P1 domain recapitulates our experimental data (e.g. Figure 6 and (13, 35))

The RMSD observed in Figure S2 is mainly a consequence of fluctuations of the 2 key loops of the P2 domain: the A’B’ (residues 289-296) and E’F’ (residues 380-384). In these simulations, when the A’B’/E’F’ loops are in the ‘closed’ conformation, the distance between them is ∼10Å with the A and B chains started in the open and closed positions, respectively, for these simulations (Figure 9). There is an apparent difference in the A’B’/E’F’ loop flexibility between the apo and the V339I mutant, despite the mutations being distal to both loops. With the wt P domain, both the A (open) and B (closed) chains took ∼600ns to move to a state with both loops in the closed conformation. In contrast, the V339I A (open) conformation did not converge to a state with the closed conformation but the closed conformations were briefly sampled during the calculation. Specifically, at ∼400 ns, A and B briefly switched conformations but then return to their original states, suggesting a facile equilibrium between the states. Interestingly, unlike wt, V339I does not form a stable structure where both A and B have the same open/closed conformation. For both wt and V339I, the C’D’ loop did not move to the “up” position and, in fact, that region was not sampled. Therefore, the C’D’ loop movement takes a much longer time than was sampled with these simulations, indicating a barrier between states or different solution conditions which are needed for that movement to be observed.

**Figure 9.**
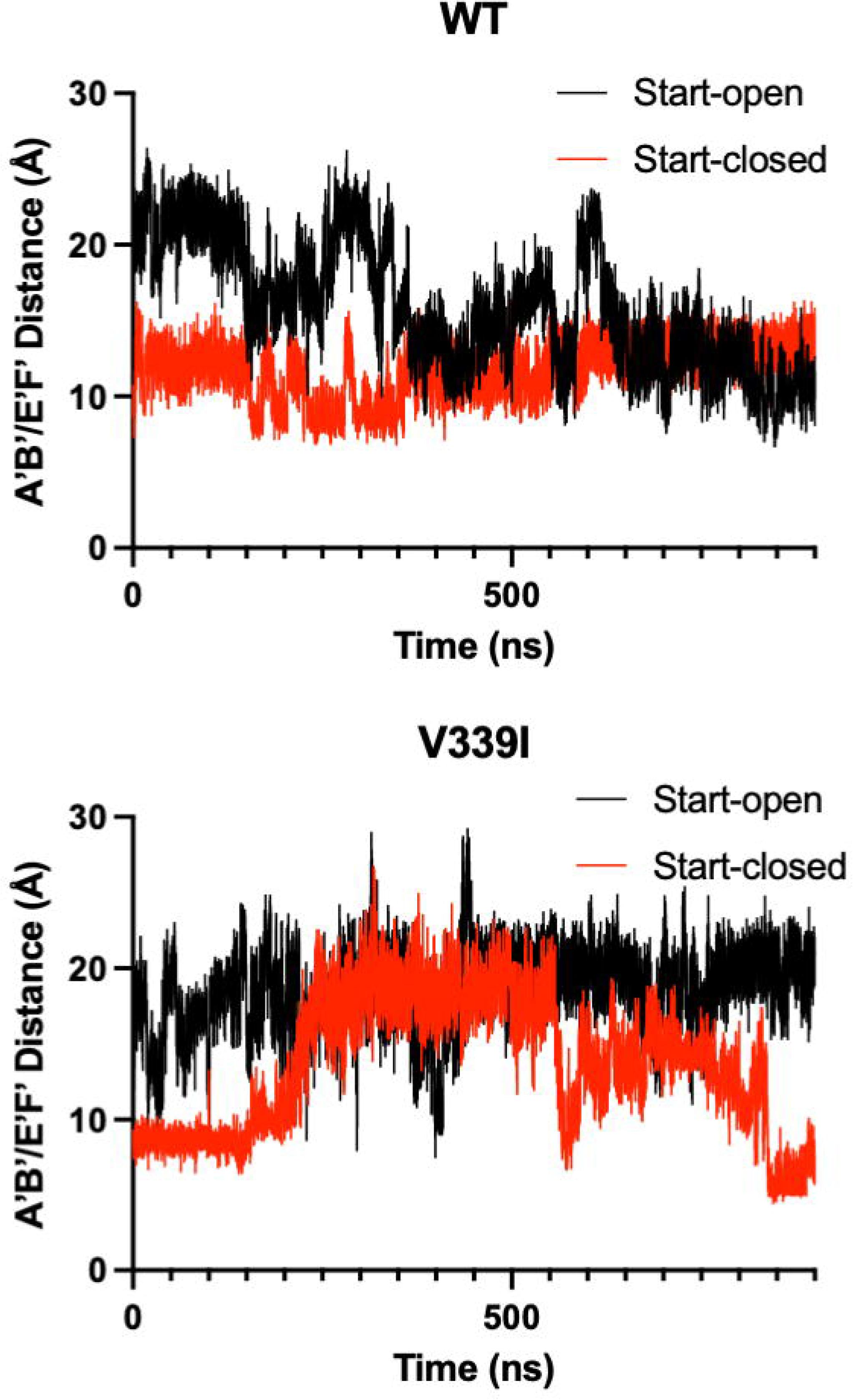
Movement in the A’B’/E’F’ loops during molecular dynamic simulations. The simulations were started with the apo crystal structure of the P domain where the A subunit (black lines) is in the ‘open’ position and the B subunit (red lines) is in the ‘closed’ position with an average distance of ∼20Å and ∼10Å, respectively. With the wt structure, the two subunits converged to approximately the closed conformation. In contrast, the V339I structure (wt with V339 replaced with an isoleucine), The V339I A subunit (open) does not converge to the closed conformation but does briefly sample an approximate conformation during the calculation. V339I appears to destabilize the conformation of the A’B’/E’F’ loops.

### P1 Movement

The radius of gyration and RMSD (Figures S1 and S2) illustrate that the P1 domains are stable and fluctuate about a mean structure during the simulation. For each chain, the P1 domains maintain conformational similarity to the original apo starting structure. However, there is a rotation of the A/B P1 domains about each other (Figure S3). To quantify this motion, the angle between the α1 helices were measured. For each chain a vector is defined by the Cα atoms of residues 457, 459, 461 and 463. The angle between those two vectors was then determined as shown in figure S3. The angle between the 2 helices fluctuates between 110° and 140° degrees with a mean value, shown by the red circle, of 125° (wt) versus 123° (V339I), with a standard deviation of 5°. A small rotation of chain B relative to chain A was similarly observed experimentally (Figure 2 (27, 30, 31, 35)). Like the radius of gyration calculations (Figure S1), the angle between the P1 domains of V339I may be slightly more fluid than wt (as per the greater deviations from the average angle) suggesting less conformational stability and more motion.

## Conclusions

The studies described herein were undertaken to understand how the V339I and D348E mutations block antibody neutralization even though they are ∼20Å from the epitope. Our previous work showed that mAbs 2D3 and 4F9 neutralized all the A6.2 escape mutants (26). Together with the fact that the escape mutants to the 2D3/4F9 antibodies were distal to the A6.2 epitope, it seemed logical at the time to assume that 2D3/4F9 bound in a different location than A6.2. However, this was dispelled with the cryo-EM structure of the 2D3 Fab’/MNV complex (13). While the orientation of the bound 2D3 was not identical to A6.2, there was extensive epitope overlap at the top of the P domain, which was distal to the locations of the escape residues V339I and D348E. Indeed, Figure 3 clearly shows that these escape mutants block all three of these antibodies. Therefore, the conformational changes caused by the V339I and D348E allosteric escape mutants disrupt the important contact points for all three antibodies. The fact that those two escape mutations did not arise in the presence of A6.2 suggests that 2D3 and 4F9 make more extensive contact and is harder to thwart. The structures of the V339I and D348E mutants demonstrate they escape all antibody neutralization by converting MNV into the activated form without the need of activators. Both appear to accomplish this by crowding regions of the C’D’ loop with larger amino acids to force it out of the apo conformation and into the ‘up’ activated position. Having V339I in the same ‘activated’ conformation in the apo, +Ca^2+^, and +GCDCA forms also allowed us to examine the effects of the activators on the critical loops that bind to antibody or receptor. While all the A’B’/E’F’ loops are all in the activated, ‘closed’, conformation, they become far less mobile when activators are added.

The molecular dynamic simulations recapitulated many of these observations and finally lend insight into the relative importance of the loop movements shown in Figure 2. In both wt and V339I simulations (Figure 9), the C’D’ loops were not observed to rotate to the ‘up’ position found in fully activated P domains as observed in the activated P domain structures (27, 30–32, 35) indicating such a change would take more than a microsecond kinetically.. Nevertheless, the A’B’/E’F’ loops were able to sample the open and closed conformations with the V339I appearing to be more fluid in these motions. Therefore, it seems likely that the C’D’ (and G’H’) loop movement controls the activation process by limiting the conformational space for the A’B’/E’F’ loops rather than imposing a structural change. Interestingly, the simulation also showed that the A and B subunits rotate about each other (Figure 2) that is necessary to facilitate the contraction of the P domains onto the shell. Again here, the V339I mutant appears to be more mobile than wt. Therefore, these studies showed that the A’B’/E’F’ loop movement and P1 domain rotation can occur spontaneously in wt virus and is accentuated by the V339I mutation. Therefore, the C’D’ loop acts more like a ‘gate keeper’ by limiting the conformational space of the A’B’/E’F’ loops rather than directly imposing a particular conformational change. By being located on the C’D’ loop, the V339I and D348E mutations both lie on the C’D’ loop and may decrease the energy required for the full activation process to occur.

This ease of conformational changes might predict the presence of intermediate virus states with mixtures of raised and retracted P domains in the virions. However, no such intermediates were observed in the raw cryo-EM data or 2D classes. This suggests that the rotation and retraction of the P domain onto the shell is a cooperative process. This cooperativity may be due to changes in the interactions between the P domain spikes in the icosahedron. When the P domains are resting on the shell, the A and C subunits make ∼260Å^2^ contact in the icosahedron (5). This connects the P domain dimers around the 5-fold axes to those at the 2-fold. As the P domains rotate and lift off the surface (apo structure), interactions switch to the P1 domains (24). Further, the rotation within the dimer during activation (19, 27, 30, 31) may switch the P1 interactions in the apo form to P2 interactions in the activated form. Therefore, these interactions may be the cooperativity that switches the P domains from one state to the other, with the activators supplying the small input of energy necessary to start the transition. In this way, activation is initiated by with the loss of P1 interactions with the pull of P2 interactions and newly formed complementarity with the shell.

Interestingly, our recent work on the MNV strain CR6 represents a case quite the opposite of these allosteric mutants. CR6 is less infectious (19) and requires multiple stimuli for full capsid activation. While V339I and D348E are nearly in the fully activated state without stimuli, CR6 requires both low pH and bile to fully activate (19). At low pH alone, the CR6 P domain has dropped onto the shell, but remains markedly disordered. This is greatly improved upon the addition of bile. Therefore, while the P domain overall has only two positions (floating above or collapsed onto the shell), the loops at the tip of the P domain have multiple conformations that become restrained upon activation.

Most viruses exist at a precipice where some environmental trigger and/or receptor starts a cascade of irreversible conformational changes initiating the infection process. Similarly, MNV responds to the low pH, and high bile and metal ion concentrations to undergo the transition to the activated state that is optimized for receptor binding. Unlike other viruses, however, this process is wholly reversible. We propose that MNV has developed this as a unique mechanism for immune evasion and these allosteric escape mutants are mimicking this process. Unlike most viruses, MNV infects gut tissue where conditions are completely different to any other location in-vivo. Rather than just surviving the trip through the alimentary canal, MNV utilizes these extreme conditions to completely alter its capsid conformation. In this way, without the need for escape mutations, it can evade the antibody response while in the gut. This model is detailed in Figure 10. Within the gut, MNV is in the activated state (red particles) and enters the epithelium via the M cells. Bile enters the small intestinal tissue via passive transport while calcium and magnesium are absorbed by the epithelium via paracellular and transcellular transport. Therefore, there are significant concentration gradients of metal ions and bile salts in the tissue directly underneath the intestinal epithelium. This allows for increased infectivity of CD300lf-expressing cells (e.g. macrophages and dendritic cells), promoting replication and shedding. As the virus and infected cells drain from the epithelium, bile and metal ions are reabsorbed and the virus reverts to the apo state. Therefore, what is presented to the lymphatic system is the apo form of MNV with its open A’B’/E’F’ loop conformation and raised P domain. In this way, the virus can repeatedly infect the gut while being effectively neutralized in the circulation. Importantly, this can explain why infection with high doses of MNV does not protect the gut from subsequent challenge but does protect the spleen where conditions keep MNV in the apo form (15). It seems likely that during the initial response to infection the high concentrations of antibodies to apo MNV might afford some protection in the gut, but it becomes prone to subsequent infection as the antibody production wanes. Added to this, bile acids are well-known to induce a tolerogenic adaptive T and B cell response (38). It is tempting to speculate that bile suppresses the generation of neutralizing antibodies to the activated form of the virus, thus exacerbating the immunological blind spot for activated MNV. Interestingly, a recent study showed that mice with higher levels of bile in the gut have concomitantly higher MNV disease scores (39), making an in-vivo connection between bile salts and infection. This dynamic and aggressive immune avoidance is only possible because of the extreme environmental differences between the gut and the circulation and the fact that MNV has an extremely flexible capsid where relatively small energy inputs can trigger complete remodeling of the capsid. Since HNoVs similarly have a contracted conformation at low pH (e.g. (1)) and are in the expanded state (apo) in PBS just like MNV (28), it will be interesting to determine how well these results translate to HNoVs. For example, some studies have suggested that adaptive immunity to HNoVs is transient wherein human volunteers infected with norovirus had symptomatic infections when challenged with the same stock 6 months later (40).

**Figure 10.**
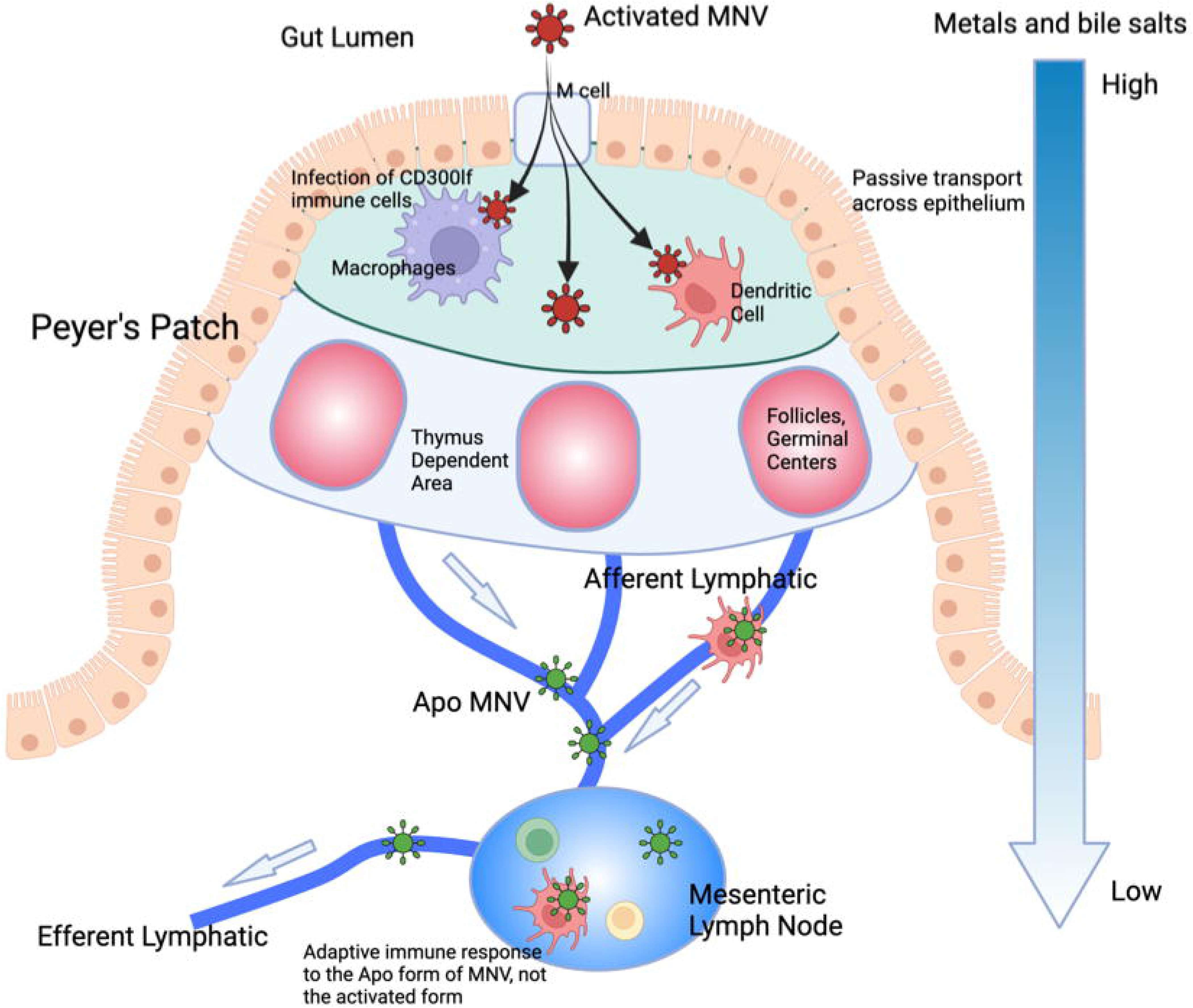
Possible function of the reversible activation process in MNV. We propose that this reversible activation may have evolved to leverage the extreme conditions in the gut to avoid immune recognition. The gut has a low pH with high concentrations of metal ions and bile salts. Each of these conditions causes the collapse of the P domain onto the shell (red virion) and buries epitopes at the tip while enhancing receptor binding (5, 30, 31). In the epithelium, passive transport of bile salts and metals allow for enhanced infection while simultaneously bile salts dampen the immune response (55). As the virus drains into the lymphatic system, those metabolites dissipate, and the virus adopts the apo structure (blue virion). The immune system therefore only sees this apo structure and therefore does not recognize the activated form in the gut. In this way, the virus can reinfect the gut in subsequent challenges without requiring escape mutations to avoid the antibody response. However, since the immune system recognizes the apo form, extraintestinal sites are protected in subsequent challenges.

Perhaps the largest potential impact of these findings is on vaccine development. Because of this rather clever means to avoid immune recognition in the gut, it is necessary to consider these potential conformational changes in the HNoV vaccine design. Theoretically, immunization with a perpetually activated form of the norovirus capsid would be anticipated to result in better protection against infection. Future studies in the MNV animal model will be important to test this concept.

## Materials and Methods

### Antibody production and purification

All antibodies were grown in 1 l Wheaton CELLine Flasks. RPMI media without serum was used in the 1 l bulk chamber and Gibco Cell MAb Medium (animal component free, Cat. 220513) that contained necessary growth factors was used for the 15 ml cell compartment. Approximately twice a week, 15 mls of cells and media were harvested from the cell compartment and replaced with MAb Medium. The cells were removed by centrifugation for 10 min at 10,000g. To the supernatant, an equal volume of saturated ammonium sulfate was added and the mixture stored at 4°C.

Monoclonal antibody A6.2 is an IgG and therefore purified using a protein G column. Since Protein G is relatively insensitive to salt concentrations, the antibody was purified directly from the ammonium sulfate precipitate. The precipitated A6.2 was centrifuged at 10,000g for 10 min and the pellet resuspended in ∼10x the volume of the pellet in PBS buffer. The sample was then loaded onto the protein G column, washed extensively with PBS buffer, and eluted with 0.1M glycine buffer, pH 2.7. The low pH of the elution was immediately neutralized using a 1M Tris buffer, pH 8.0. If not used immediately, the antibody was again precipitated with ammonium sulfate.

Monoclonal antibodies 4F9 and 2D3 are IgA antibodies and therefore could not be purified with the protein G column. For these, the precipitated mixtures from the CELLine flasks were centrifuged and resuspended in PBS as above. Approximately 10mls of this resuspension was loaded onto an Cytiva XK-50 column (300mm x 50mm i.d.) filled with Sephacryl S-300 media. Fractions containing antibody were pooled and stored in ammonium sulfate. The precipitate was collected by centrifugation (10,000g for 10 min) and dialyzed against 25mM Tris buffer, pH 7.4. The protein was then purified using a Mono-Q 5/50 GL anion exchange column on an AKTA system. Buffer A was the same as used for dialysis while buffer B had an additional 1 M NaCl. Using a gradient of 1% buffer B/ml, the protein eluted at 0.1-0.15 M NaCl. The antibody fractions were pooled and stored as an ammonium sulfate precipitate.

Purity for all antibodies were monitored with SDS-PAGE. Immediately before use, purified antibodies were harvested from the ammonium sulfate precipitate and dialyzed against PBS. Concentrations were determined using an extinction coefficient at 280nm of 1 mg/ml*OD. For the experiments comparing relative efficacy, the antibody concentrations were normalized to each other by analyzing the SDS-PAGE using ImageJ (41) and integrating the heavy chain bands.

### Virus production and purification

The V339I and D348E mutants of MNV-1 were essentially produced and purified as previously described (5). In brief, BV-2 (RRID:CVCL_0182) cells were grown in spinner 4 l spinner flasks in Media B (Gibco S-MEM, 0.1% [wt/v] Kolliphore P188, 1% vol/vol NEAA, 0.062g/l Penicillin, 0.4g/l Streptomycin sulfate, and 26mM sodium bicarbonate) until they reached a density of ∼0.5-1.0×10^6^ cells/ml. ∼6 liters of cells were harvested by centrifuging 4,000g for 10 min. The cells were suspended in 1 liter HEPES (AH) media, placed into a 4l flask, and ∼1×10^9^ pfu’s of MNV was added. Importantly, we found that both mutations reverted to wt after several passages as we scaled up production. To prevent this, ∼10µg/ml of purified 2D3 was added to the infection. The suspension was transferred to a 37° incubator without CO_2_ and shaken at ∼70 rpm in the dark. Mutant replication was apparently slower than wt in that the infected cells required 48 hours of incubation for maximal titer compared to 24 hrs for wt. The infected cell suspension was centrifuged for 30 min at 5,000g and the supernatant was collected. To the supernatant, dry NaCl and PEG 8,000 was added to yield 0.3M and 10%, respectively. The solution was then mixed at 4°C overnight. The solution was then centrifuged for 30 min at 5,000g and the pellet was resuspended in 50-80 mls of PBS. After the suspension was incubated at 4° C for several hours, the debris was removed by centrifugation at 10,000g for 30min. To the supernatant, glycerol was added to a final concentration of 10% (v/v) as a cryoprotectant, divided into 1.5ml aliquots, and stored at −80°C.

Immediately before the cryo-EM experiments, 50-100 mls of this material was thawed and centrifuged at 45,000 rpm for two hours in a 50.2 Ti Beckman rotor. The pellets were resuspended in less than a total of 3 mls of PBS and allowed to incubate for several hours at 4°C. Debris was removed by centrifugation and the supernatant was then layered onto Beckman SW41 tubes containing 7.5-45% linear sucrose gradients using PBS as buffer. After centrifugation for 1.5-2.0 hours at 35,000 rpm at 4°C, MNV forms a band ∼2/3 of the way down the tube. The virus was collected via puncturing the side of the tube with a syringe. The pooled bands were dialyzed overnight against buffer containing 30 mM HEPES, pH 7.4, and 100mM NaCl. When the titer of the frozen virus was less than 10^7^ pfu/ml, the virus band was often difficult to see. A large section of the gradient near the expected location was collected, the virus pelleted as above, and the sample reseparated on a sucrose gradient. The act of pelleting forces some of the cellular contaminants into a pellet that could not be resuspended. A portion of the virus was used to confirm the presence of the mutation using RT-PCR and Sanger sequencing as previously described (42). The virus was pelleted using a Beckman 50.2 Ti rotor for 1.5 hours at 45,000 rpm at 4°C and resuspended in the same buffer. The virus was divided into three different portions to which nothing, 1 mM CaCl_2_ (final concentration), or 10 mM GCDCA (final concentration) was added. The samples were immediately frozen for cryo-EM studies.

### Antibody neutralization assays

Virus neutralization plaque assays were performed as previously described (27). BV2 cells from the spinner cultures were added to 6-well tissue culture plates (9.5 cm^2^ surface area) and allowed to attach for ∼2 hours. The excess media was removed and 0.5 mls of virus diluted to ∼10^3^-10^8^ pfu/ml with Media B. For neutralization, purified monoclonal A6.2 was added to a final concentration of ∼50µg/ml and incubated with the virus for ∼1 hr. To each well, 0.5 mls of diluted virus (+/- antibody) was added and the virus was allowed to attach for 45min on a tilting shaker. After incubation, the media was removed and 2 mls of 50% (v/v) low melting temp agar/P6 media (31mM BSA, 79mM magnesium chloride hexahydrate, Gibco MEM, 2% [v/v] NEAA, 0.124g/l Penicillin, 0.8g/l Streptomycin sulfate, and 52mM sodium bicarbonate) was added to the wells. Once the agar was solidified, 5 mls of Media B was added to each well. The plates were incubated at 37°C and 5% CO_2_ for 37 hours. After incubation, the media and agar were removed, and a 0.1% crystal violet/20% ethanol mixture was added for plaque visualization. Since the various virus samples had different titers, the neutralization efficacy was calculated by the number of plaques without antibody divided by the plaque count with antibody added.

### Cryo-electron microscopy (cryo-EM)

MNV samples were at concentrations of ∼1 mg/ml. The virus was vitrified as previously described (43) on carbon holey film (R2×1 Quantifoil®; Micro Tools GmbH, Jena, Germany) grids. Briefly, grids were cleaned in Gatan 950 Solarus plasma cleaner for 40 s in hydrogen-oxygen gas mixture. 4 µl of the V339I or D348E virus solutions were applied to the holey films, blotted with filter paper, and plunged into liquid ethane. The EM-GP2® (Leica) automated plunger was used for vitrification.

The grids were screened for ice and sample quality and imaged in a Titan Krios G3i (Thermo-Fisher) microscope. The microscope was equipped with a BioQuantum electron energy filter (Ametek, Inc.) and operated at 300 keV. A slit width of 20 eV was used for data collection. Images were acquired with SerialEM (44) using fast acquisition mode with beam-image shift used for hole centering instead of stage movement. The direct detector camera (K3, Ametek) operated in counted mode, images were recorded with overall electron dose of 40 electrons/ Å^2^; the defocus range was −1.5 to −2.5 µm. The pixel size was 1.096 Å on the specimen scale.

The data collection statistics are summarized in Table 1.

**Table 1:**
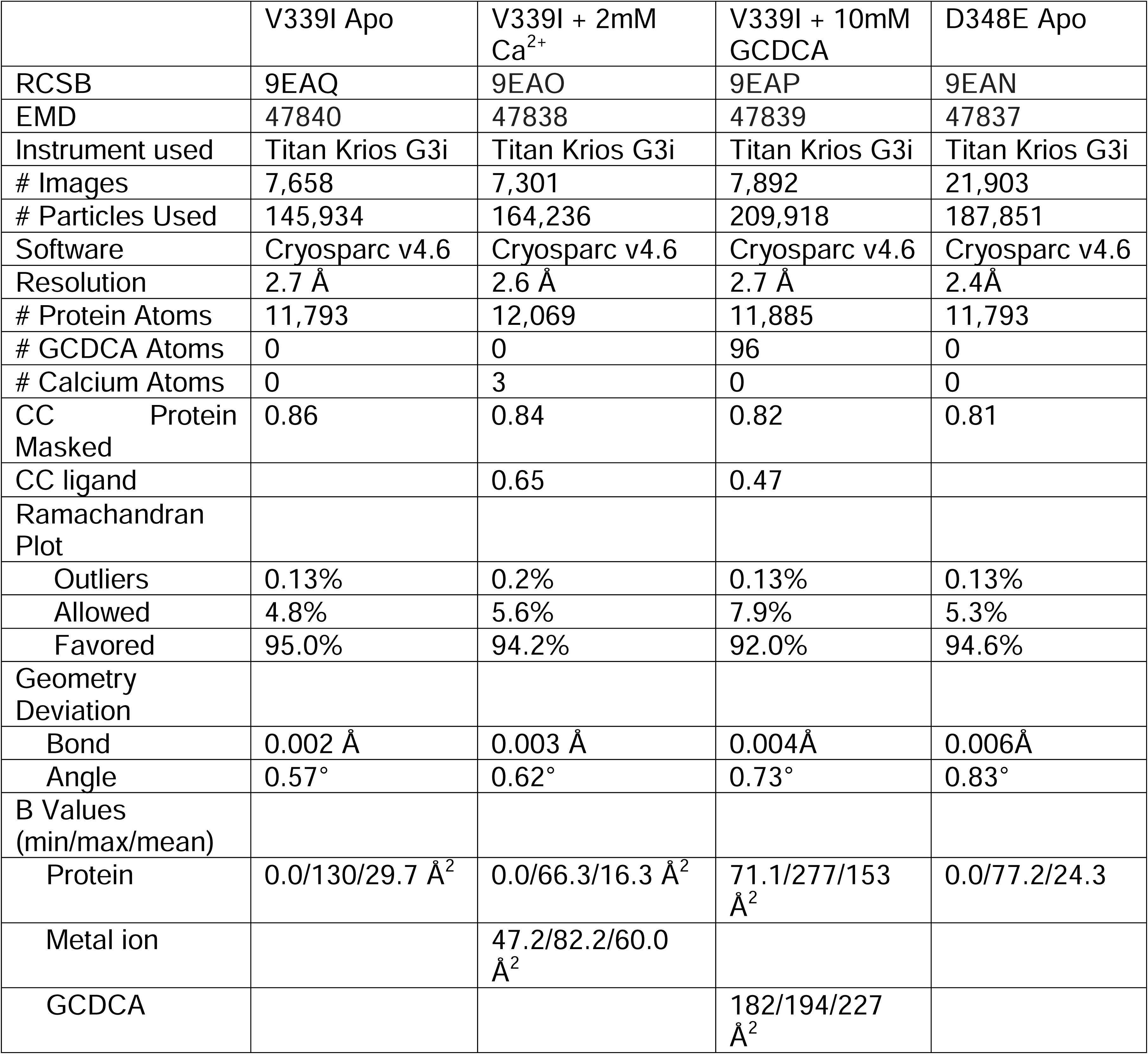
Data and refinement statistics for the four structures.

|  | V339I Apo | V339I + 2mM Ca<sup>2+</sup> | V339I + 10mM GCDCA | D348E Apo |
| --- | --- | --- | --- | --- |
| RCSB | 9EAQ | 9EAO | 9EAP | 9EAN |
| EMD | 47840 | 47838 | 47839 | 47837 |
| Instrument used | Titan Krios G3i | Titan Krios G3i | Titan Krios G3i | Titan Krios G3i |
| # Images | 7,658 | 7,301 | 7,892 | 21,903 |
| # Particles Used | 145,934 | 164,236 | 209,918 | 187,851 |
| Software | Cryosparc v4.6 | Cryosparc v4.6 | Cryosparc v4.6 | Cryosparc v4.6 |
| Resolution | 2.7 Å | 2.6 Å | 2.7 Å | 2.4 Å |
| # Protein Atoms | 11,793 | 12,069 | 11,885 | 11,793 |
| # GCDCA Atoms | 0 | 0 | 96 | 0 |
| # Calcium Atoms | 0 | 3 | 0 | 0 |
| CC Protein Masked | 0.86 | 0.84 | 0.82 | 0.81 |
| CC ligand |  | 0.65 | 0.47 |  |
| Ramachandran Plot |  |  |  |  |
| Outliers | 0.13% | 0.2% | 0.13% | 0.13% |
| Allowed | 4.8% | 5.6% | 7.9% | 5.3% |
| Favored | 95.0% | 94.2% | 92.0% | 94.6% |
| Geometry Deviation |  |  |  |  |
| Bond | 0.002 Å | 0.003 Å | 0.004 Å | 0.006 Å |
| Angle | 0.57° | 0.62° | 0.73° | 0.83° |
| B Values (min/max/mean) |  |  |  |  |
| Protein | 0.0/130/29.7 Å<sup>2</sup> | 0.0/66.3/16.3 Å<sup>2</sup> | 71.1/277/153 Å<sup>2</sup> | 0.0/77.2/24.3 |
| Metal ion |  | 47.2/82.2/60.0 Å<sup>2</sup> |  |  |
| GCDCA |  |  | 182/194/227 Å<sup>2</sup> |  |

### Image processing

For the MNV reconstructions, 7,658, 7,301, 7,892 and 21,903 images were used for the apo V339I, V339I+calcium, V339I+GCDCA, and apo D348E samples that yielded 145,934, 164,236, 209,918, and 187,851 particles, respectively. Our model from our previous image reconstructions (e.g. (27, 31)) was used to calculate 25 templates and particles were picked using the Cryosparc 4.6 (45) template picker. Particles were culled using 2D classification. A low pass filter was applied to our previous structure and the non-uniform refinement algorithm yielded a cryo-EM density with Gold Standard Fourier shell resolution of ∼2.7Å for all three V339I reconstructions and 2.4Å for the D348E reconstruction.

### Structure Refinement

All structure refinement was performed using PHENIX (46). The density of all four reconstructions had the activated conformation where the P domain had contracted onto the shell as previously observed in the MNV/bile complex (5) and therefore that structure was used as an initial model for building and refinement in PHENIX. To ensure that the model of the icosahedral asymmetric unit did not move into density of adjacent subunits, the model used for real space refinement included neighboring subunits that formed the A/B and C/C dimers. The model was manually fitted into the EM envelope and several cycles of real space refinement (rigid body, global minimization, and simulated annealing) in PHENIX (46) and rebuilding in COOT (47) were performed. The final refinement statistics are shown in Table 1.

### Molecular Dynamics Simulations

The initial structure used for the MD simulation was the x-ray structure of the apo P domain, PDBID: 3LQ6, with a resolution of 2.0Å. This structure has an asymmetric conformation with the A’B’/E’F’ loops of chain A in the “open” conformation and chain B in the “closed” conformation (12). The V339I model was modeled using the Mutate option in VMD (48). Hydrogen atoms were added, using a protonation state of pH 7.0, and each chain was capped with acetylated N-termini and N-methylamidated C-termini using PSFGEN (48). The system was then solvated with TIP3P water with at least 20 Å between the protein and the box edge. The system was neutralized and NaCl ions were added for a salt concentration of 0.11M.

The molecular dynamics simulations were performed with version 2.14 of NAMD (49) with the CHARMM36m force field parameters (50). Initially, the energy was minimized using the conjugate gradient algorithm for 20K steps then heated slowly for 25K steps from 0K to 300K with no restraints. Next, the protein atoms were constrained, and the solvent and ions were allowed to equilibrate for 3 ns in the isothermal-isobaric (NPT) ensemble. The constraints on the protein atoms were subsequentially removed and all the atoms were allowed to move for an additional 5 ns of equilibration. A further 900 ns was simulated using the isothermal-isobaric (NPT) ensemble at a temperature of 300 K and 1 atm. A 2 fs timestep is used to integrate the equations of motion for the 900 ns of production. Particle mesh Ewald (51) was used to calculate the long-range electrostatic interactions and van der Waals interactions were truncated at 12 Å. All bonds were constrained with the RATTLE algorithm (52). Trajectory frames were saved every 2 ps.

Analyses of the simulation data were performed with CPPTRAJ (53), VMD (48), and PyMol (54) for the residues of the P domain only, i.e., residues 237-522 of each chain. For the root-mean-square deviation (RMSD) calculations, each snapshot was first aligned to the first frame after equilibration, considering only the protein backbone atoms. RMSD values were then calculated for all atoms except for hydrogens for each chain, A and B, and each domain, P1 (residues 416-522) and P2 (residues 237-415). Mass-weighted radius of gyration (R_g_) was calculated for all non-hydrogen atoms of each domain, P1 and P2.

## Acknowledgments

This work was supported by an NIH grants 1R01-AI141465 (TJS), R21-AI154647 (CEW), R01-GM037657 (BMP), and a McLaughlin Fellowship (FC). The authors would like to acknowledge the support of the Sealy Center for Structural Biology at UTMB. The coordinates have been deposited to the Protein Data Base with accession codes of 9EAQ, 9EAO, 9EAP, and 9EAN for V339I apo, V339I calcium, V339I GCDCA, and D348E apo structures, respectively. The related cryo-EM maps have been deposited with the EMDB accession codes of 47840, 47838, 47839, and 47837, respectively. The authors acknowledge the Texas Advanced Computing Center (TACC) at the University of Texas at Austin for providing the computational resources used for the molecular dynamics simulations reported within this paper. URL:https/www.tacc.utexas.edu

