## Supplemental Figure 1 for "Murine norovirus allosteric escape mutants mimic gut activation"

Figure S1

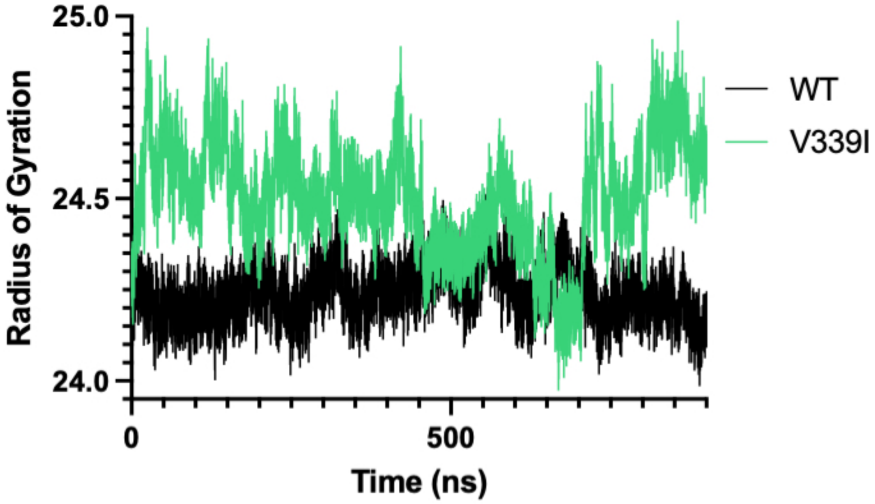


Figure S1: The radius of gyration of wt (black) and V339I (green) during the 900ns simulation. While both structures were stable during the simulation, V339I appeared to be more mobile.
