## Supplemental Figure 2 for "Murine norovirus allosteric escape mutants mimic gut activation"

Figure S2

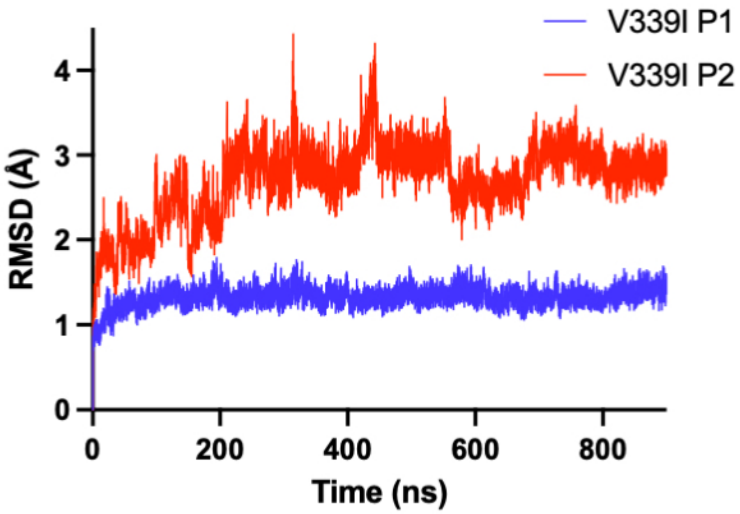


Figure S2: Root-mean-square deviations of apo V339I separated according P1 (blue) and P2 (red) domains. Note that the P2 domain is far more mobile than the P1 domain.
