## Supplemental Figure 3 for "Murine norovirus allosteric escape mutants mimic gut activation"

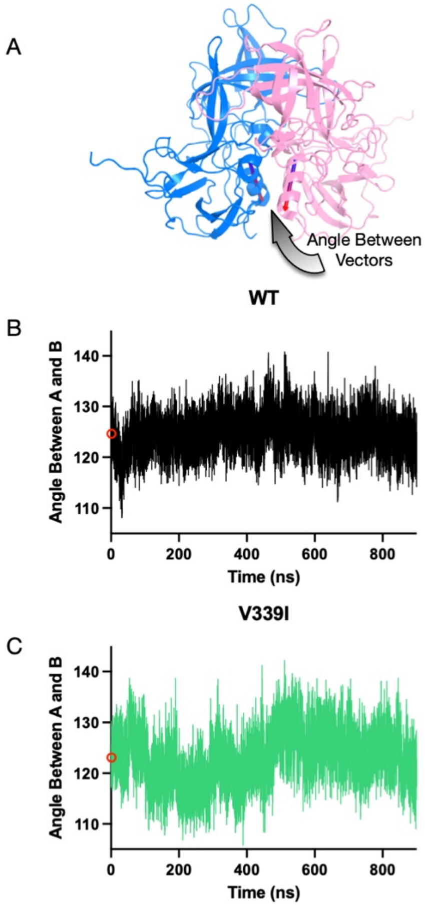


Figure S3. Rotation of the A and B subunit P1 domains during molecular dynamic simulations. A) To measure the angles between the two domains (blue and pink), vectors were calculated using a helix in the P1 domain (arrow). B,C) During the simulations the angle between the A and B P1 domains sampled a range of values like that observed in previous cryo-EM and crystal structures (Figure 1). The red circle denotes the average angle during simulation. While the average angles of wt and V339I are within the margin of error, V339I appears to be more fluid than wt.
